# Trajectory Classification Method for Anchored Molecular Motor-Biopolymer Interactions in the C. elegans first Mitosis

**DOI:** 10.1101/2022.09.11.507465

**Authors:** John Linehan, Alan Edwards, Vincent Boudreau, Paul Maddox

## Abstract

During zygotic mitosis, forces generated at the cell cortex are required for the separation and migration of paternally provided centrosomes, pronuclear migration, proper segregation of genetic material, and successful cell division. Identification of individual cortical force generating units *in vivo* is necessary to study the regulation of microtubule dependent force generation throughout the cell cycle, to further understanding of asymmetric cell division, and to identify the molecular mechanism of force generation. Here we present a method to determine both the location and relative number of microtubule dependent cortical force generating units using single molecule imaging of fluorescently labelled dynein. Dynein behavior is modeled to differentiate and categorize trajectories that correspond to that which is cortically bound and interacting with a microtubule, and is cortically bound and not interacting with a microtubule. The categorization strategy recapitulates well known force asymmetries in the first mitosis of the C. elegans embryo. To evaluate the robustness of categorization, we RNAi depleted the microtubule subunit TBA-2 resulting in reduction of the number of trajectories categorized as engaged with a microtubule. This technique will be a valuable tool to provide new insight to the molecular mechanisms of dynein cortical force generation and its regulation as well as other instances wherein anchored motors interact with biopolymers (eg. Actin, tubulin, DNA).

## Introduction

In the first embryonic division of the *C. elegans* embryo, polarity is established with entry of the paternal nuclei and associated centrosome during fertilization (1, 2)(3, 4). At the culmination of zygotic mitosis, this asymmetry leads to the division of the embryo into two differently sized daughter cells, with distinct cell fates ((5)) ((6)). During zygotic prophase, microtubules emanating from the duplicated centrosomes interact with the cell cortex resulting in centrosome separation, pronuclear migration, and asymmetric positioning of the mitotic spindle and cleavage furrow (7). A cortical force generating complex, consisting of a pair of redundant heterotrimeric G alpha subunits (GOA-1 and GPA-16) bound to the plasma membrane, the go-loco protein GPR1/2 and Numa like protein LIN-5, enables microtubule dependent force generation (8) (9)(10, 11). In this complex, dynein binds directly with LIN-5, anchoring it to the cortex to allow its motor domain to interact with microtubule plus ends emanating from the centrosomes (12)(13). Once bound to the force generation complex, dynein either: a) anchors microtubule plus ends in place so that force generation may occur through microtubule depolymerization, b) pulls on microtubules by motoring toward the minus ends anchored to the centrosomes, or c) a combination of both (14) (15, 16) (17). Prior work used confocal microscopy of fluorescently labeled plasma membrane to detect small invaginations as a proxy to identify the number of cortical force generating units (18). Alternatively, locations of force generators have been identified by studying microtubule dynamics at the cortex, revealing two populations of microtubules associated with active pulling and pushing forces respectively (19).

In contrast, we look to categorize cortically located dynein trajectories collected using single molecule imaging with the 25^th^ and 75^th^ percentile values of step length within a trajectory. Here, we used Total Internal Reflection Fluorescence (TIRF) imaging to detect single dynein molecules at the zygote cortex. Subsecond imaging rates and super-resolution positioning of the signal generated trajectories that encode force generation events. To decode trajectories, we developed the classification method presented here. Our technique enables the identification and localization of cortically bound dynein that is interacting with a microtubule, distinguishing it from the cortically bound population that is not.

Fertilization of the embryo also cues the onset of cortical flows, redistributing cytoplasmic proteins throughout the embryo (reviewed in (20), (1)) simultaneous to centrosome separation and migration due to cortical forces translated by astral microtubules (21). In prophase, the ratio of LIN-5 fluorescence intensity in the anterior relative to the posterior extremes is biased towards the anterior by roughly 1.5x (9). The ratio of GPR1/2 in the anterior relative to the posterior extremes of the embryo is roughly 1.35 (9). This data suggests that in prophase, the anterior cortex contains a larger number of locations where dynein can cortically bind and generate force on microtubule plus ends.(12).

Spindle positioning is a visual output of how force producing elements are arrayed upon the embryo cortex. During anaphase, the spindle is just posterior to the midplane, indicating that the force equilibrium has shifted, likely due to redistribution of the dynein based force producing units (22) (23). Confirming this, optically induced centrosome disintegration experiments show that astral fragments of the spindle located posteriorly travel faster in the direction of the posterior cortex than anteriorly located fragments move in the direction of the anterior cortex (24). Additionally, the ratio of LIN-5 fluorescence intensity in the anterior end during anaphase relative to the posterior is roughly 0.93 (9) and that of GPR1/2 roughly 0.8 (9). These data indirectly show where and when dynein functions to deliver force to the mitotic apparatus. In this work, we aim to characterize dynein activity as it binds to and acts on the microtubule plus end.

Here we present a model of cortically bound dynein that includes both microtubule bound and unbound states. Using this model we run a Monte-Carlo simulation that characterizes the step length distribution in the (x,y) plane of both behaviors. We then use these distributions to categorize cortically located dynein trajectories in the anterior and posterior during the *C. elegans* first embryonic division to determine whether the model characterization effectively recapitulates the well know asymmetry in force generation at these two stages of mitosis. We then test the model against a depletion of the microtubule subunit TBA-2, effectively reducing the number of microtubules in the cell during prophase. Having categorized trajectories, we present a distribution of the time dynein is bound to the force generation complex and thus able to contribute to the movement of centrosomes to set up asymmetric cell division.

## Results

We first considered the *in situ* context in which microtubule-dependent cortical force generation occurs in the *C. elegans* embryo. Single Molecule imaging performed using Total Internal Reflection Fluorescence Microscopy (TIRFm) resulted in numerous dynein trajectories at the cortex (see methods) (25). Dynein is bound to the force generation complex consisting of membrane bound G alpha, GPR1/2, and LIN-5 (12, 22). In our model, this structure acts as a lever that extends normal to the membrane. The lever moves both horizontally and vertically about the point where G alpha is bound to the cortex, so that the force generation complex and dynein swivels about a fixed point in space. This swiveling about the fixed point constrains dynein to move along the surface of a hemisphere when bound to the force generation complex (Figure 1C, E).

**Figure 1:**
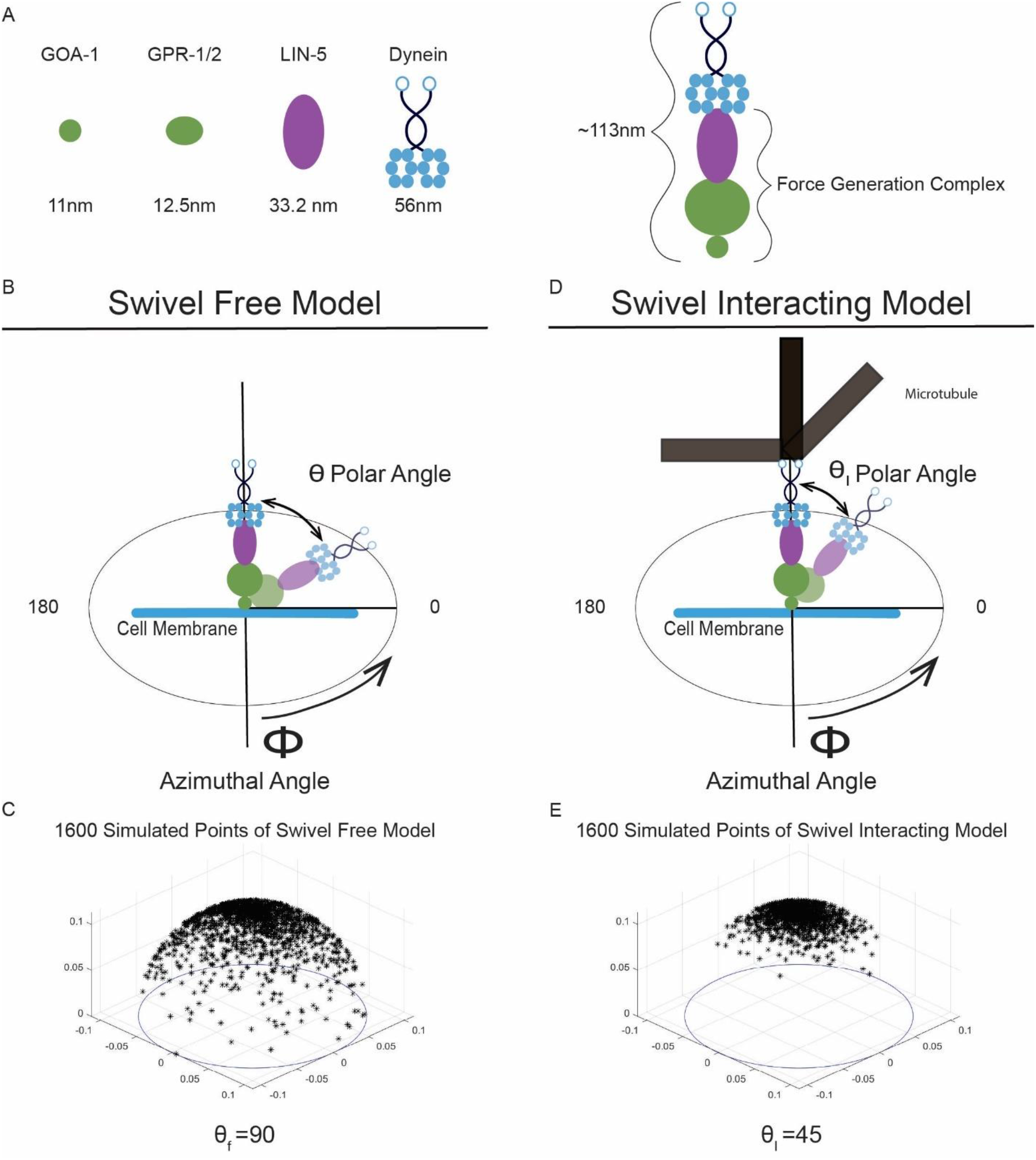
Force Generation Complex Description, Size, and Dynein model. A) Proteins making up the force generation complex and relative size, full force generation complex and dynein length. B) Swivel Free model, dynein and force generation complex deflect off the line normal to the membrane. C) Simulation of Swivel Free Model. D) Swivel Interacting model, depiction of force generation complex and dynein interacting with a microtubule. E) Simulation of Swivel Interacting model.

Stochastic forces act to swing the lever arm about the point where G alpha is bound to the membrane. We assume that the likelihood of deflection from the line normal to the membrane follows a Gaussian distribution, so that larger deflections from the normal line are less likely to occur. Dynein and the force generation complex reach into the dense filamentous actin cortex patchwork to interact with a microtubule, where the density of the filamentous actin likely constrains the force generation complex and dynein to this orientation (26) (27). RNAi of the non-muscle motor protein NMY-2 in the first mitosis of the *C. elegans* embryo resulted in an increased number of membrane invaginations protruding into the cytoplasm (18), leading to the conclusion that the actin cortex plays a role in constraining the force generation complex and dynein’s position.

We refer to the angle that the lever arm swings out from the line normal to the membrane as the polar angle. A new polar angle is randomly selected for each iteration of the simulation. The most likely value for the polar angle in our simulation is 0 degrees, so that it is most often standing normal to the membrane. The polar angle theta at each iteration is given by *θ*(*t*) = *N*_*θ*_(*µ*, 0.34*θ*_*max*_), where *θ*_*max*_ is the maximum degree of deflection from the line normal to the membrane.

Movement in the x-y plane parallel to the membrane is given by a uniform distribution. The angle swept out in the (x,y) plane is called the azimuthal angle (Figure 1B,D). The azimuthal angle is also randomly selected for each time step of the simulation so that *ϕ*(*t*) = *U*(−*π, π*).

The globular protein G alpha is the smallest protein in this complex with an estimated diameter of 10.95nm (Figure 1A). GPR1/2 consists of two subunits, GPR-1 and GPR-2, that are modeled here as having a total diameter of 12.45nm (Figure 1A). LIN-5 is a coiled coil protein that binds to GPR1/2 and dynein, it is estimated to be 33.24nm long (Figure 1A). Dynein length is estimated to be 55.92nm (Figure 1A; see materials and methods for protein radius and length estimates). In total the force generation complex from tip to tail, consisting of G alpha, GPR1/2, LIN-5, and dynein, standing normal to the membrane is 113nm (Figure 1A). This is the length of the lever arm normal to the membrane.

We assume that the force generation complex stays intact throughout the time that dynein is present, so that the structure is not turning over while dynein is bound to the force generation complex consisting of G alpha, GPR1/2, and LIN-5. We use a dynein heavy chain labelled with a green fluorescent protein (eGFP). The enhanced green fluorescent protein has a length of 4.2nm and width 2.4 nm (28). The fluorescent label is sufficiently close, less than 4.2nm away, to the dynein heavy chain so that the fluorescent label provides an accurate readout of dynein location including consideration of the resolution limit of Total Internal Reflection Imaging (see materials and methods).

The location of the GFP tagged dynein when bound to the cortex through the force generation complex is modelled using a spherical coordinate system with position coordinates (*ρ, θ, ϕ*). *ρ* is the radial coordinate, defined as the tip to tail length of the force generation complex and dynein, *ρ* = 0.113 *µm. θ* is the polar angle that measures the degree of deflection from the line normal to the cell membrane, and *ϕ* the azimuthal angle that describes position in the (x, y) plane. The force generation complex is bound at the location where G alpha binds to the membrane, and is free to swing across a full circle in the azimuthal angle *ϕ ∈* [([−*π, π*])]. The position of the fluorescent tag is given by *r*(*ρ, ϕ, θ*).

After binding a microtubule, the polar angle is constrained so that the force generation complex bound to dynein cannot deflect further from the line normal to the membrane than it can in the absence of a microtubule. Dynein processivity on a microtubule acts as a constraint that limits the degree of deflection towards the membrane. We refer to this model as the swivel model. Dynein that is cortically bound through the force generation complex and interacting with a microtubule is called the *swivel interacting* group (Figure 1D, E). Dynein that is bound to the force generation complex but **not** interacting with a microtubule is the *swivel free* group (Figure 1 B, C). Simulation data for the Swivel free group (Figure 1 C) and swivel interacting group (Figure 1 E) are shown to detail the impact on step length of two different maximum polar angles of deflection. The spherical coordinates generated from simulation are mapped to the Cartesian coordinate system with the transform equations,

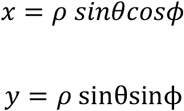

to match the output from dynein trajectories collected using total internal reflection fluorescence microcopy (29). The position of the fluorescent tag is then given by,

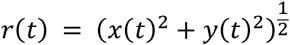

Where r is the position of the fluorescent molecule over the course of the trajectory. We characterize the swivel model’s behavior using the step length distribution generated from the simulation. The step lengths within a trajectory are calculated as,

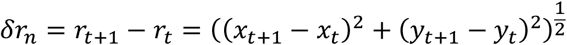

Where n is the number of steps within a trajectory, the length of the trajectory minus one point.

### Determining the Maximum Polar Angle of Deflection for Swivel Free and Swivel Interacting Models that Best Fit Measured Step Length Probability Mass Function

The swivel model differentiates between cortically bound dynein that is interacting with a microtubule, and cortically bound dynein that is not interacting with a microtubule; by specifying the maximum polar angle, defined to be the degree of deflection from the line normal to the membrane, of both groups (Figure 1 B,D). The maximum value for the polar angle is determined by Monte-Carlo simulation.

Given a specified polar angle in the *swivel free* group, the *swivel interacting* model was evaluated for each angle from 10 degrees up to the specified swivel free maximum polar angle, in increments of 5-degrees with one hundred iterations. The simulated swivel models’ step length distributions were concatenated, and the probability mass function of step length was calculated. The normalized root mean sum of squared error is used to determine the pair of maximum polar angels of deflection that best recapitulated the probability mass function of step length from the measured dynein trajectories (Figure 2A). Cortically located dynein trajectories were collected TIRF Imaging (see materials and methods).

**Figure 2:**
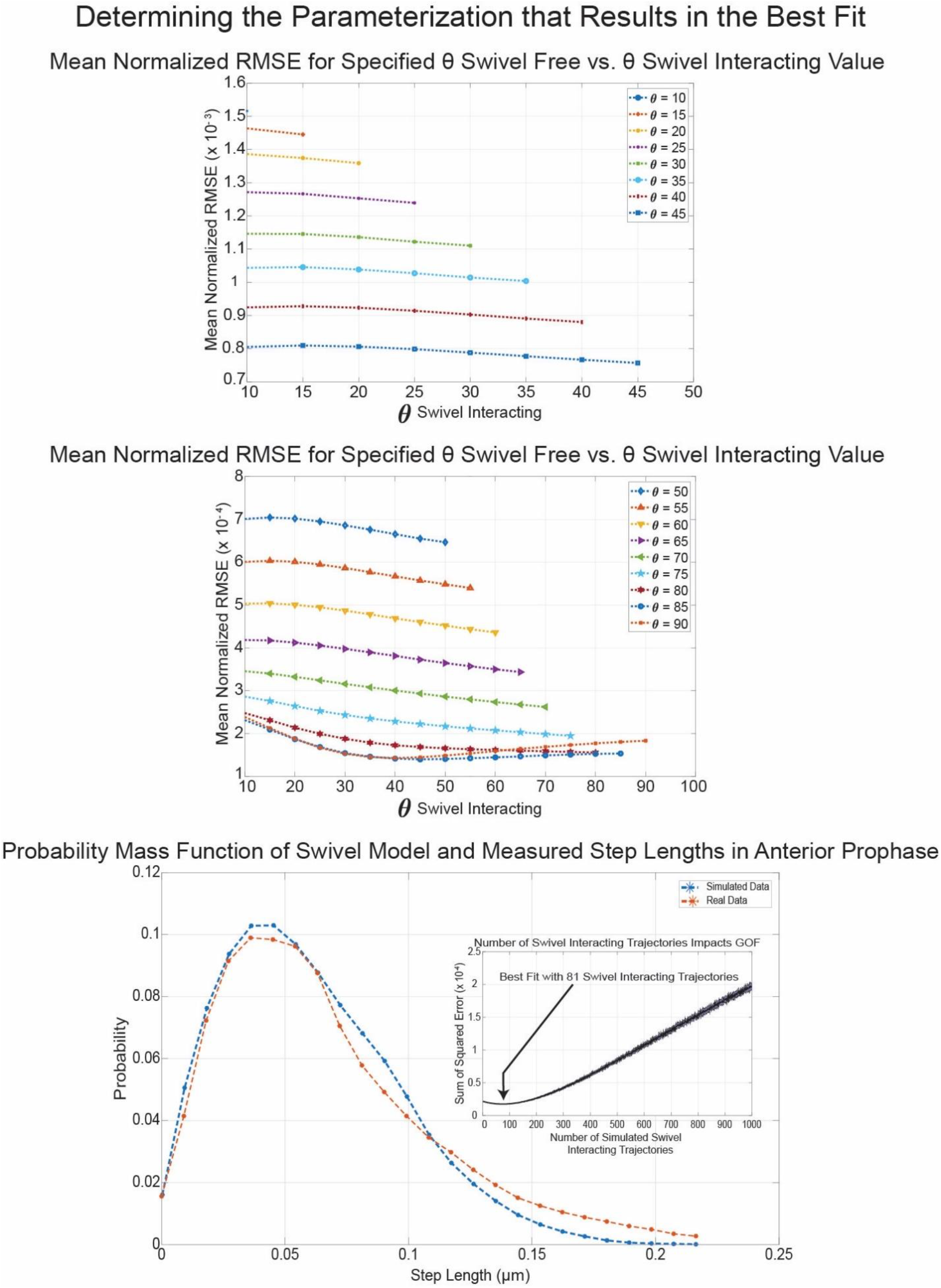
Determining the Parametrization that Best Fits Measured Step Length Probability Mass Function: A) Evaluation of maximum polar angle of deflection for swivel free model from 10-45 degrees. B) Evaluation of maximum polar angle of deflection for swivel interacting model from 50-90 degrees. C) Comparison of Simulated probability mass function of step length for optimal polar angles of deflection. Sub Panel C) Sum of Square Error of Fit when varying the number of simulated swivel interacting trajectories. The number of swivel interacting trajectories affects the likelihood of short steps.

The parametrization with the smallest normalized average root mean square error is when the max deflection angle is *θ*_*f*_ = 85^⋅^ for the swivel free model, and *θ*_*I*_ = 45^⋅^ for the swivel interacting model (Figure 2B blue line and circles). This parameterization has normalized root mean square error of 1.4*x*10^−4^ (Figure 2B, C). The measured step length probability mass function in the anterior end of the zygote does extend out to a step length of 0.5 *µm*, however the probability of a step occurring that is 0.25*µm* is near zero (Figure 2B). The fit of the simulated probability mass function depends on the number of simulated swivel interacting trajectories (Figure 2C subpanel). We ran distinct simulations with the specified maximum angles of deflection for the swivel free and swivel interacting groups (Figure 2 A,B), iterating over the number of swivel interacting trajectories simulated. We found that the simulation fit could be improved using a concatenation of 1000 swivel free trajectories and 81 swivel interacting trajectories (Figure 2C). The sum of square error decreases up to 81 trajectories, and then increases in value up to 1000 trajectories (Figure 2C subpanel). That fewer swivel interacting trajectories result in a better fit likely indicates that there are more dynein in the *swivel free* group than in the *swivel interacting* group. A high density of cortically bound dynein is likely necessary to ensure that astral microtubules are sufficiently likely to encounter a force generation complex.

### Swivel Model Step Length Characterizations and Likelihood of Miss Categorizing a Trajectory

The step length probability mass function of both swivel free and swivel interacting groups is determined by Monte-Carlo Simulation (30). The median step length of the swivel free group is 0.0575*µm*, 25^th^ percentile step length is 0.0352*µm*, and the 75^th^ percentile value is 0.0851*µm* (Figure 3A). For the Swivel interacting group the median value is 0.0316*µm*, 25^th^ percentile value is 0.0191*µm*, the 75^th^ percentile value is 0.0471*µm* (Figure 3 B). The 25^th^ and 75^th^ percentile values are used as the categorization metric to classify a trajectory as belonging to either the swivel free or swivel interacting groups. The 75^th^ percentile of the swivel interacting group overlaps with the 25^th^ percentile of the swivel free group (Figure 3C).

**Figure 3:**
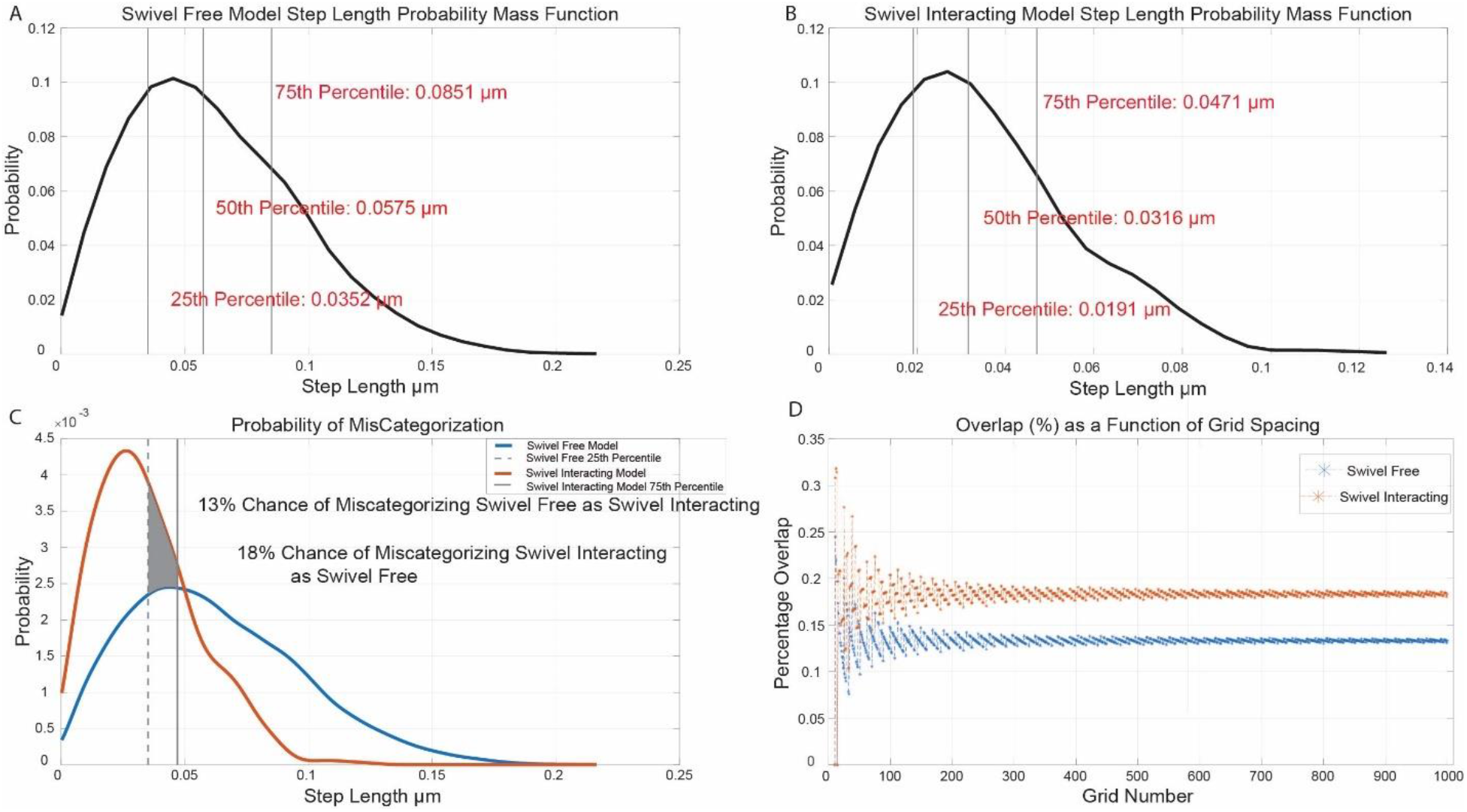
Swivel Model Step Length Distributions and likelihood of mis-categorization. A) Probability of step length versus step length for swivel free model simulation, with 25^th^, 50^th^, and 75^th^ percentile step lengths marked. B) Swivel Interacting Model Step length probability vs. Step Length, with 25^th^, 50^th^, and 75^th^ percentiles marked. C) Overlay of Swivel Interacting and Swivel Free Models, probability versus step length. The shaded region indicates the overlap of the 25^th^ percentile value of swivel free model and 75^th^ percentile of swivel interacting model. D) Percentage Overlap of swivel free and swivel interacting showing that the overlap does not vary significantly with alteration of “bin size”, discretization of space between maximum and minimum steps.

The overlap of the two probability step length curves was calculated to determine the percentage of miss categorizations one can expect using the swivel model as a classification tool. Using these criteria, we expect that 13% of the swivel free group will likely be improperly classified as swivel interacting trajectories and 18% of the swivel interacting trajectories will likely be improperly classified as swivel free trajectories (Figure 3C). To determine that the likelihood of mis categorization was not impacted by the binning or grouping of values to calculate the probability mass function, we created a probability density function using a kernel filter for both models. We then evaluated both model groups probability functions over unique discretization’s of the space between the maximum and minimum value of displacement simulated. We found that the overlap of the two probability mass functions does not vary greatly with the discretization of the support for a number of grid points greater than 100 (Figure 3D).

We ran simulations of dynein undergoing Brownian diffusion near the cortex using a step length distribution generated using a previously reported value for cytoplasmic dynein diffusion (31). The median step length for dynein undergoing Brownian motion is 0.22 *µm*, while the 25^th^ and 75^th^ percentile values, used for categorization, are 0.14 *µm* and 0.31 *µm*. These values are ten-fold greater than expected for either of the swivel models. The likelihood of classifying a dynein molecule that is undergoing Brownian diffusion near the cortex as being bound to the cortex is significantly low so that it is ignored.

### Application of the Swivel Model to the Categorization of Dynein Trajectories Collected in the Anterior and Posterior end of the C. elegans first embryonic division during Prophase and Anaphase

After fertilization in the C. elegans zygote, paternally donated centrosomes coordinate centrosome and pronuclear migration by cortically generated forces transmitted through microtubules (22, 32). This event is followed by the meeting of maternal and paternal nuclei in the midplane, which is again coordinated by the microtubule cytoskeleton from forces generated at the cortex (22) (Figure 4A). During prophase the anterior end of the zygote is known to have a greater amount of the force generation components LIN-5 and GPR1/2 than the posterior, making it likely that the anterior contributes more force to this process than the posterior end, resulting in asymmetric cell division critical for cell fate determination in development (9).

**Figure 4:**
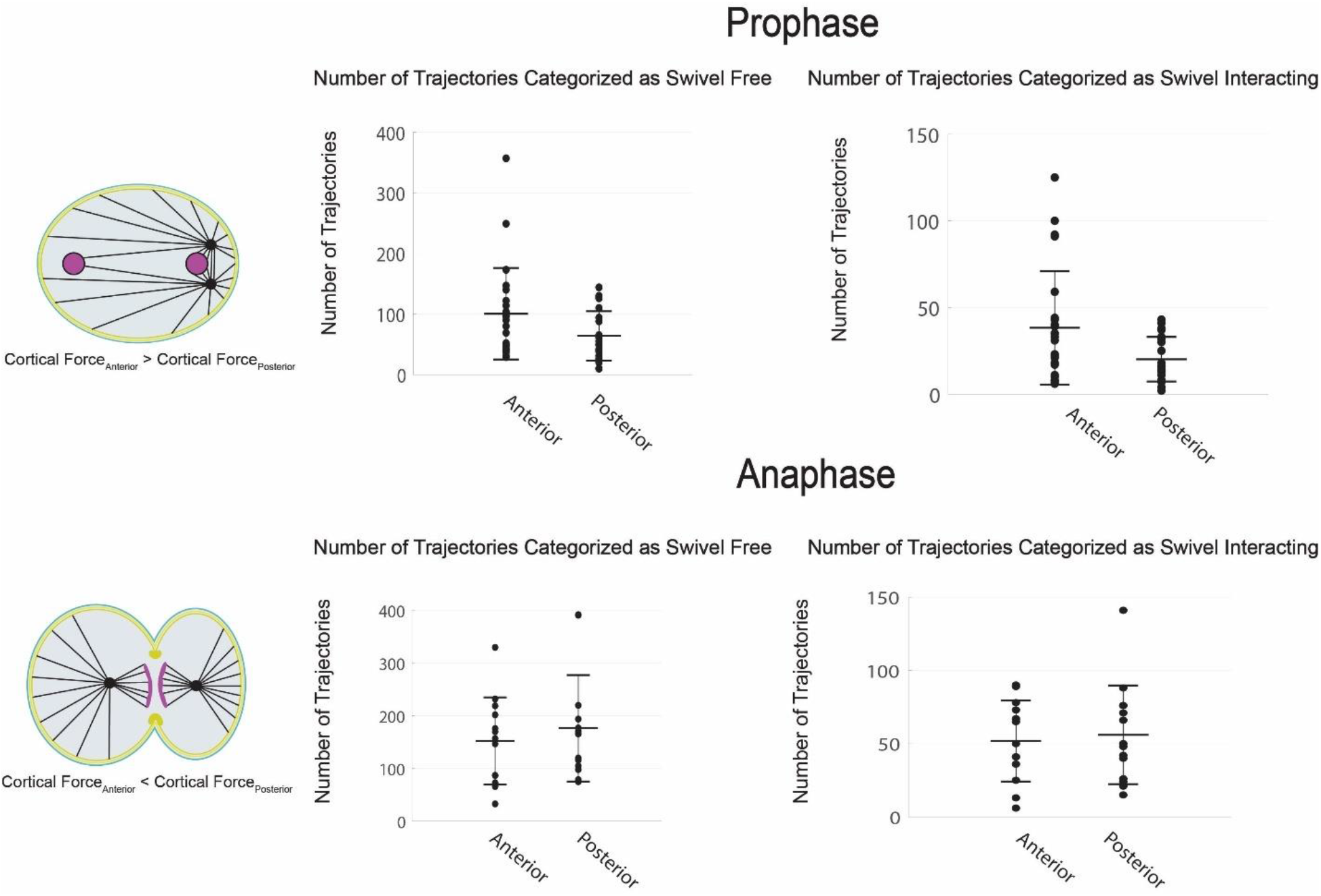
Comparison of Number of Trajectories Categorized as Swivel Free and Swivel Interacting. A) Schematic of the embryo during prophase. B) Prophase Zygote comparison of trajectories categorized as swivel free in posterior and anterior of the embryo. C) Comparison of the number of trajectories categorized as swivel interacting during prophase in the anterior and posterior. D) Schematic of the embryo during anaphase. E) Comparison of the number of trajectories categorized as Swivel Free during anaphase in the anterior and posterior of the zygote. F) Comparison of the number of trajectories categorized as swivel Interacting during anaphase in the anterior and posterior of the embryo.

We used Total Internal Reflection Fluorescence microscopy to image the anterior and posterior ends of the C. elegans zygote during prophase using a C. elegans strain with dynein heavy chain tagged with eGFP (33). Images were collected every 0.027 seconds over the course of one minute, with a pixel size of 0.06*µm x* 0.06*µm*. The displacement length between dynein locations within a trajectory were determined using Gaussian super positioning software freely available through FIJI (34)(35). The median of each trajectory’s step length was used to categorize that trajectory as belonging to the swivel free or swivel interacting group. In each zygote the total number of both groups were tabulated. In the anterior end of the zygote during prophase the number of dynein trajectories categorized as swivel free is 100 ± 75 trajectories per minute (Figure 4B)(n = 24). In the posterior end during prophase the number of trajectories categorized as swivel free is 64 ± 41 trajectories per minute (Figure 4B) (n = 23). In the anterior end of the embryo during prophase 38 ± 33 trajectories per minute were categorized as belonging to the swivel interacting group (Figure 4C). In the posterior end of the embryo 20 ± 13 trajectories per minute were classified as belonging to the swivel interacting group (Figure 4C). The number of classified trajectories recapitulate the well-known force asymmetry during prophase of the first embryonic division in C. elegans.

We next tested the categorization method against trajectories collected during anaphase. In anaphase the spindle is positioned slightly closer to the posterior cortex (22) (Figure 4D). Optically induced centrosome disintegration experiments have shown that when the spindle is ablated anteriorly located spindle fragments move towards the anterior cortex slightly slower than posterior fragments move towards the posterior cortex (24). The fluorescence intensity of LIN-5 and GPR1/2 also indicate an increased number of force generating units in the posterior in comparison to the anterior (9). Although the total force generated in the posterior of the embryo is greater than the total force generated in the anterior, the difference in magnitude is slight. Categorizing dynein trajectories during anaphase is a challenge to the methods ability to properly discriminate trajectories given the slight difference in force asymmetry at this stage.

In the anterior end of the embryo during anaphase 152 ± 83 trajectories were classified as belonging to the swivel free group over the course of one minute (Figure 4 E). In the posterior 176 ± 34 trajectories per minute were categorized as being in the swivel free group (Figure 4E). In the anterior end during anaphase 52 ± 28 trajectories per minute were classified as swivel interacting (Figure 4F). In the posterior end during anaphase 56 ± 34 trajectories per minute were classified as belonging to the swivel interacting group. Here the mean values of the swivel free group reflect the previously identified asymmetry in the amount of force generating units in the embryo during anaphase.

The large standard deviation present across groups motivated us to perform a bootstrapping experiment to determine the likelihood of categorizing the number of trajectories presented in Figure 4 due to chance (Supplemental figure 1). We found that the distribution of the number of trajectories classified was statistically significantly different in each of the above cell cycle phases and locations in the embryo for both models (supplemental figure 1), indicating that the method is categorizing based of a biological mechanism rather than by chance.

### Testing Swivel Model Categorization Method against tba-2 (RNAi) depletion and Characterizing Dynein Cortical Binding Time

We tested the swivel model based categorization scheme against depletion of the microtubule subunit alpha tubulin 2 (TBA-2). Microtubules are made up of dimers consisting of alpha and beta tubulin proteins (36). We performed a 16 hour depletion of the alpha tubulin protein (expected 30% reduction in protein, longer depletions lead to pleotropic defects precluding analysis). In the anterior end of tba-2 (RNAi) depleted embryos during prophase there were 14 ± 9 trajectories per minute categorized as swivel interacting (Figure 5A). In the posterior end there were 16 ± 9 trajectories per minute classified as swivel interacting (Figure 5B). In total we see a 52% reduction in the mean number of trajectories categorized as swivel interacting when comparing prophase embryos to tba-2 (RNAi) depleted embryos (Figure 5B). We performed a bootstrapping analysis to determine the likelihood that a trajectory would be classified as belonging to either model due to chance (Supplemental Figure 2). We found that the distribution of trajectories collected in both the anterior and posterior were statistically significantly different from the distribution of trajectories classified using the bootstrapping method.

**Figure 5:**
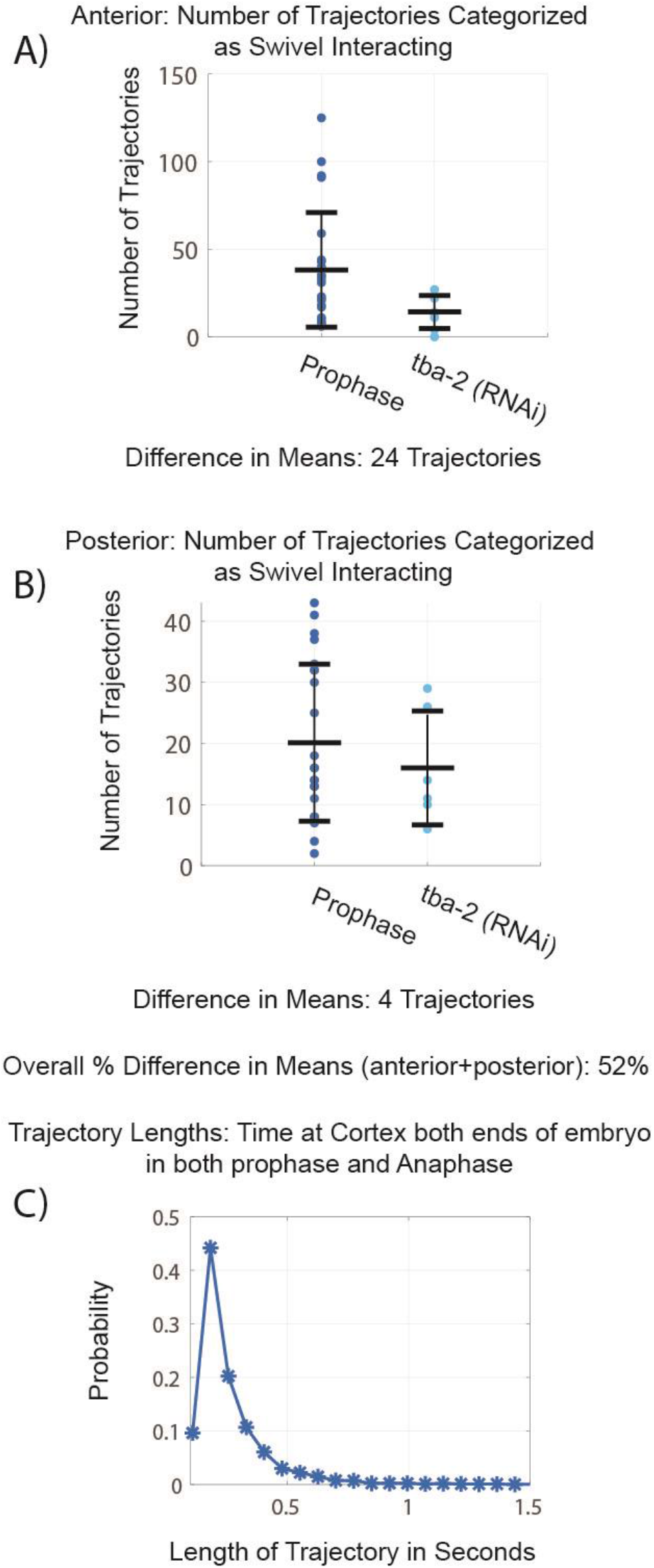
Comparing tba-2 rnai depletion data to prophase data. A) Comparison of the number of trajectories categorized as swivel interacting in the anterior end of the embryo during prophase in a tba-2 depleted embryo. B) Comparison of the number of trajectories categorized as swivel interacting in the posterior of the embryo in a tba-2 depleted embryo. C) Proportion of trajectories vs trajectory length, distribution of trajectory length, measure of time dynein is bound to the cortex.

Having tested the swivel model based categorization method against the well-known force asymmetries in prophase and anaphase of the first embryonic division in C. elegans, as well as against depletion of the microtubules; we sought to quantify the time dynein spends bound to the force generation complex and membrane by characterizing lengths of trajectories categorized in both model groups. We tabulated the number of points in each trajectory classified in both the swivel free and swivel interacting group in both ends of the embryo in prophase and anaphase (Figure 5C). The 25^th^ percentile of this distribution is 0.16 seconds, the 50^th^ percentile is 0.19 seconds, and the 75^th^ percentile is 0.27 seconds (Figure 5C).

## Discussion

We have presented a model to describe the behavior of cortically bound dynein in two unique circumstances. The first is in the absence of a microtubule, what we refer to as the swivel free group, and the second is in the presence of a microtubule, the swivel interacting group. In our model, interaction with microtubules reduces the degree from which the force generation complex and dynein can deflect from a line normal to the cell membrane. In strong support of this model, we collected high-spatial temporal images of cortically located dynein using TIRF Imaging. We determined the maximum angle of deflection for both swivel free and swivel interacting groups by simulating trajectories and fitting the model generated probability mass functions to real data. The best fitting parameterizations indicate that cortically bound dynein protrudes perpendicular to the membrane into the actin cortex. Stochastic forces act on cortically bound dynein to reposition it, but we speculate that the force generation complex generates an increased restoring force with degree of deflection, making it increasingly more difficult to deflect towards the membrane. When interacting with a microtubule we speculate that the force generation complex and dynein move about the surface of a hemisphere as dynein successively binds and unbinds from the microtubule in a background of stochastic forces. Dynein affinity for the microtubule acts to constrain the degree of deflection from normal in addition to any restoring force coming from the force generation complex.

Upon application of the model to the categorization of cortically located dynein trajectories we have found that the model classification metrics recapitulate previously identified asymmetries in force generation during prophase (9, 15, 18, 24). Where a roughly two-fold increase in both swivel free, and swivel interacting, trajectories were collected in the anterior relative to the posterior.

In anaphase the categorization technique identified a larger number of both swivel free and swivel interacting trajectories than in prophase. The model as well identified a larger number of swivel free trajectories in the posterior of the embryo than in the anterior. The number of trajectories categorized as being engaged with a microtubule, the swivel interacting group, is the same between ends of the embryo. The difference in the magnitude of force being generated from the two ends of the embryo is slight (24), however we do identify an increased number of dynein bound to force generation complexes in the posterior end of the embryo, in total there are more force generating units in the posterior than the anterior during anaphase, which of course agrees with previous biological observations (9, 15, 18, 24). We find that density of force generating units is strikingly similar between the two ends of the embryo during anaphase.

We next sought to challenge our model based categorization method by depleting the microtubule subunit alpha tubulin 2 (tba-2). An 8 hour depletion was performed that results in a 30% reduction in protein since higher levels of depletion are lethal. Since microtubules are made up of dimers of alpha and beta tubulin subunits, we expect that this depletion would have great effect in reducing the number of microtubules present in the embryo during prophase, reducing the number of trajectories classified as swivel interacting (36). We hypothesized that the swivel models accuracy could be determined by applying its categorization to tba-2(RNAi) data. We found an approximately 46% reduction in the average number of trajectories categorized as swivel interacting. We feel that the model parameterization describing cortically bound dynein microtubule interactions is indeed sensitive enough to not only identify cortically bound dynein, but to distinguish cortically bound dynein that is interacting with a microtubule from dynein that is not.

Having distinguished cortically bound dynein from cortically diffusing dynein, we next looked to determine the amount of time dynein spends bound to the force generation complex. We found that 91% of trajectories categorized as either swivel free or swivel interacting were present in our imaging at the cortex for less than .51 seconds. In comparison previous studies have used membrane invagination lifetime to quantify dynein binding time to 1 second. Roughly 9% of our trajectories were at the cortex for more than .51 seconds. (18) were likely observing this extreme end of the distribution, those dynein molecules that were bound to the membrane long enough to allow a significant protrusion to form. These findings mirror previously identified microtubule lifetimes at the cortex as well indicative of the relationship between cortical dynein binding time and microtubule dwelling time at the cortex (19).

Here we present a model of cortically bound dynein behavior that recapitulates the well-known force asymmetries in the first embryonic division of the c. elegans embryo. The purpose of this swivel model is to identify cortically bound dynein that is engaged with a microtubule so that biological study can be performed of this population, without being muddied by cortically diffusing dynein, or cortically bound dynein that is not engaged with a microtubule. As well the model draws out material properties of the force generation complex, that it stands largely perpendicular to the membrane and that it likely generates a nonlinear restoring force as it is compressed. We feel this model will be of use to researchers from numerous fields using single molecule localization microcopy methods, in addition to the study of asymmetric cell division, regulation of microtubule dependent cortical force generation throughout the cell cycle, and the study of embedded molecular motor-biopolymer interactions.

## Methods

### C. elegans use, RNAi, and microscopy

The worm strain SV1803 (he264[eGFP::dhc-1]) was grown and maintained at 20°C using standard procedures. Bacterial strains containing a vector expressing double-stranded RNA under the isopropyl β-D-1-thiogalactopyranoside promoter were obtained from the Ahringer library (Bob Goldstein’s laboratory, University of North Carolina at Chapel Hill, Chapel Hill, NC). Targets were confirmed by sequencing.

Embryos were mounted in egg buffer between a 1.5 coverslip and a 4.5% agarose pad in egg buffer and a microscope slide and sealed with VALAP (37). Embryos were imaged on a Nikon TIRF microscope with a 1.5x magnifier using a 100× Apo TIRF oil-immersion objective (Nikon), an Andor iXon3 EMCCD camera and NIS-Elements (Nikon) at 22°C. Image pixel size is 0.6 *µm x* 0.6*µm* with one image taken every 0.027 seconds. We bleached the cell for the first 1000 timepoints without quantifying dynein behavior. This was due to the high density of dynein particles at the cortex making identification of individual trajectories difficult. Our approach allowed us to have a high enough signal-to-noise ratio to track individual dynein particles.

### Image Analysis

All image analysis was done using the TrackMate (38, 39) plugin in FIJI (35). Tracking was performed using a Difference of Gaussian (DoG) detection segmenter and simple Linear Assignment Problem (LAP) tracking. Image processing was performed using a custom FIJI plug-in that is available upon request.

### Numerical Programming

All numerical programming was performed using Matlab version R2018b.

### Protein size Calculation

All protein sizes were calculated using AlphaFold, PDB, and Mol* (40–43). GOA-1, GPR1/2, and LIN-5 lengths were calculated from *C. elegans* purified EM structures. Heavy-chain dynein (*H. sapien*) with BICD-2 (*M. musculus*) and dynactin (*S. scrofa*) were used to calculate length of the force generation complex. Protein length was measured by greatest linear distance between polar amino acids.

## Acknowledgements

We greatly thank Tony Perdue (Microscopy Core, Biology Department-retired, UNC–Chapel Hill) for support. We also thank members of the Amy Maddox lab (UNC—Chapel Hill) and Paul Maddox lab (UNC—Chapel Hill), as well as Sebastian Fürthauer (TU-Wien), and Katie Newhall (UNC-Chapel Hill) for critical reading and discussion of the manuscript. This study was supported by the National Science Foundation CAREER Award 1652512 to P.S.M. J.B.L. was supported in part by a grant from NIGMS under award T32 GM119999, as well as NIH R01-102390 and NSF 1616661.

## Author contributions

A.E. and V.B designed and carried out experiments in *C. elegans* embryos and performed image and data analysis. J.B.L. designed model, and wrote MatLab based code used for data calculations, statistics, and computational modeling. J.B.L, A.E., V.B. and PM wrote the manuscript.

## Supplemental Figures – Likelihood of Trajectory Categorization Due to Chance

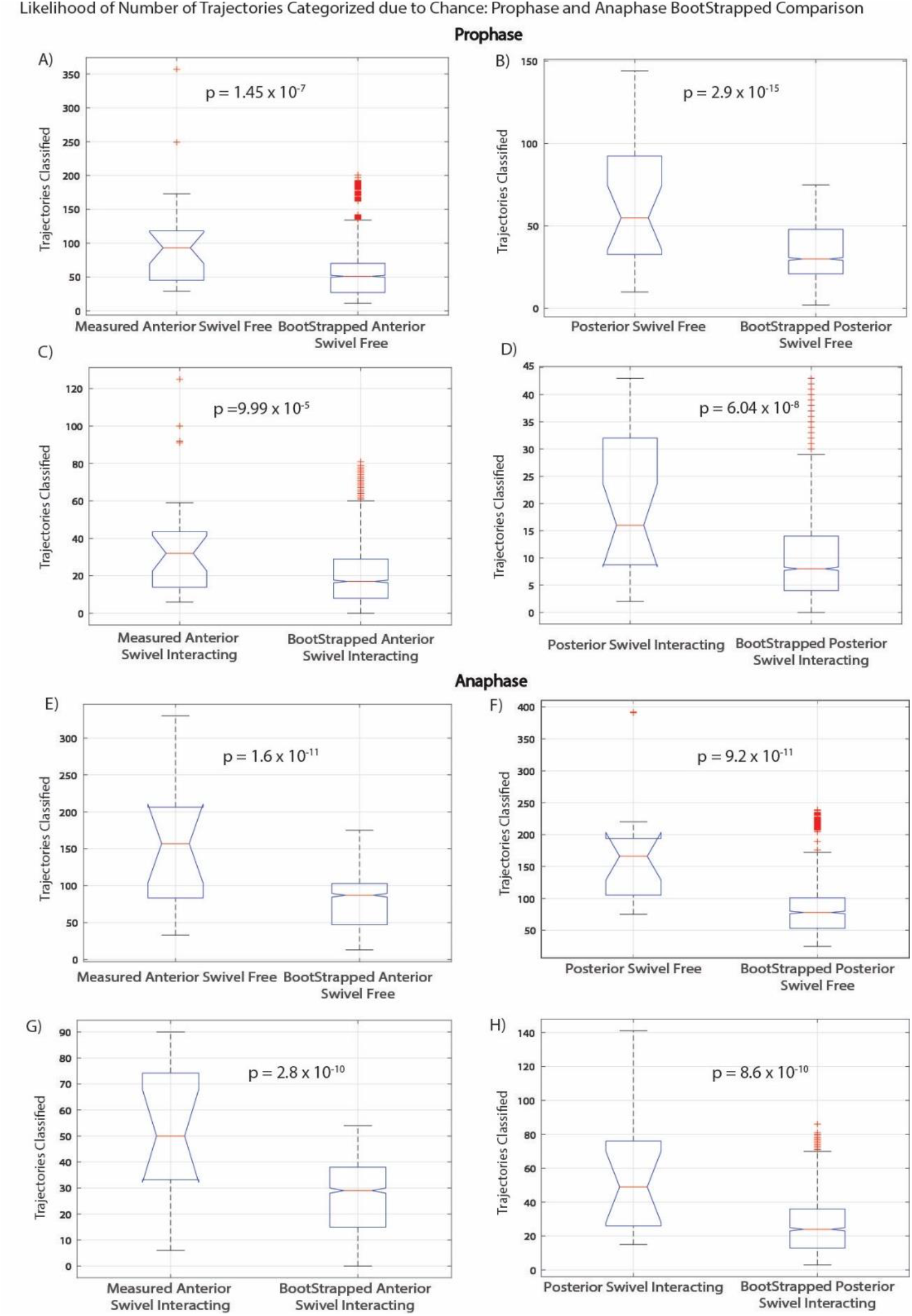

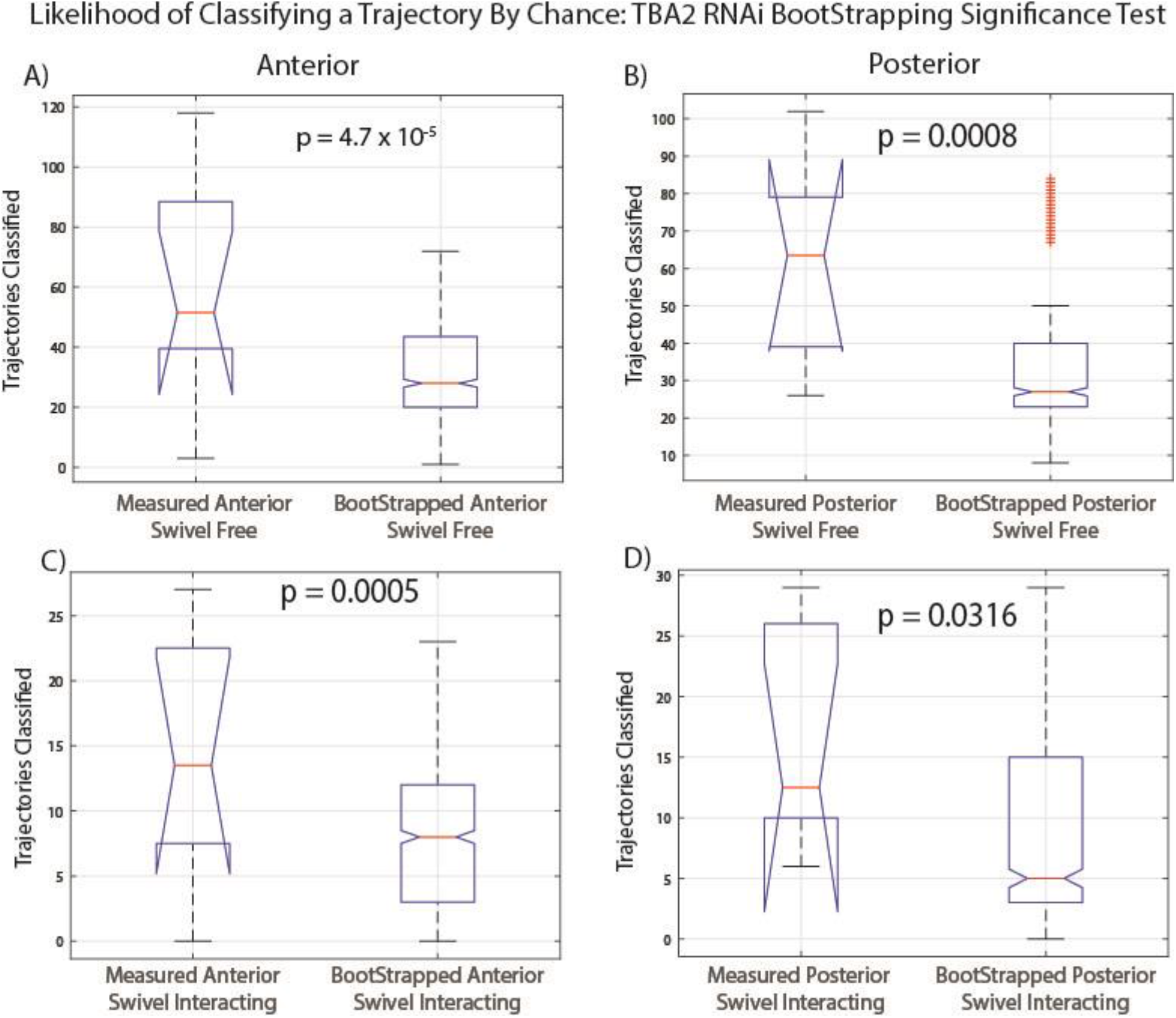

